# Coral monitoring with environmental DNA captures community restructuring after mass bleaching mortality

**DOI:** 10.64898/2026.11.24.769030

**Authors:** Luke Thomas, Defne A. Sahin, Nicole M. Ryan, Anna K. Cresswell, Carolina Castro-Sanguino, Martina Zucchi, Christy Ka Wing Kong, Matthew J. Birt, Camille M. Grimaldi, James P. Gilmour

**Affiliations:** Australian Institute of Marine Science, Indian Ocean Marine Research Centre, Crawley, WA, Australia; Oceans Institute, University of Western Australia, Perth, WA, Australia, 6009

## Abstract

Marine heatwaves are restructuring coral reefs, but monitoring coral community change across impacted habitats and reefs remains expensive and logistically challenging. Metabarcoding of seawater DNA represents a potential solution to the issue of scale, but technical challenges and a lack of ground truthing with conventional measures of coral community structure have limited its application. In early 2025, Mermaid Reef in the Rowley Shoals, a remote oceanic atoll in northwest Australia, experienced unprecedented heat stress resulting in widespread coral bleaching and mortality. Using photo-transect data alongside seawater eDNA samples, we quantified changes in coral cover and community assemblage before and after the heatwave and compared results of the two monitoring methods. Photo-transect data showed sharp reductions in live coral cover, particularly in *Acropora*, the dominant pre-bleaching genus. eDNA data mirrored these shifts, and *Acropora* sequences dominated pre-bleaching samples but declined strongly after bleaching, while several less abundant genera increased in relative read abundance. Across sites and timepoints, eDNA reads corresponded to photo-transect cover, and multivariate assemblage patterns strongly aligned between the two methods. These results demonstrate that seawater eDNA metabarcoding can provide rapid detection of community restructuring on coral reefs and can augment existing monitoring frameworks.

## Introduction

Mass coral bleaching, driven by prolonged marine heatwaves, is now the primary cause of coral mortality globally (Eakin et al., 2018, 2026; Hoegh-Guldberg et al., 2007; Hughes et al., 2017; Pandolfi et al., 2003). Monitoring community-level change following these disturbances at ecologically relevant scales remains a central challenge for coral reef management. Conventional monitoring programs rely on diver-based surveys, most commonly through methods such as manta tows or photo-transects, to quantify coral cover and community composition (Jonker et al., 2008). However, these approaches are labour-intensive, spatially constrained, and limited by depth, visibility, and access (Beijbom et al., 2015; Cresswell et al., 2021; Pearman et al., 2016). Additionally, coral reefs are often in remote locations, with limited time and researchers on site to conduct surveys.

Environmental DNA (eDNA) offers a complementary approach to community monitoring that can be deployed rapidly and repeatedly at scale to generate taxonomically resolved biodiversity data (Barnes & Turner, 2016; Elbrecht et al., 2018; Evans et al., 2017; Ip et al., 2023; Noda et al., 2025; Peiris et al., 2024; Stat et al., 2017; Thomsen & Willerslev, 2015). Despite its potential, the uptake of eDNA metabarcoding into routine monitoring frameworks has been limited, mostly due to the uncertainties associated with sequence count data and the links to more conventional metrics of community condition, such as cover, biomass, abundance, or catch per unit effort. Sequence counts derived from metabarcoding environmental samples are shaped by a combination of interacting factors, including taxon-specific shedding and decay rates, hydrodynamic transport, and PCR amplification bias, that may decouple sequence count from organism abundance (Brandão-Dias et al., 2025; Brasseale et al., 2025; Gold et al., 2023; Kelly et al., 2019; Lamb et al., 2019; Pastor Rollan et al., 2024; Shaffer et al., 2025).

This is particularly true for universal metabarcoding assays, such as the widely used 18S rRNA (Hadziavdic et al., 2014) or Cytochrome c oxidase subunit 1 (CO1-Kelly et al., 2017), that by design amplify a diverse set of taxa and thus introduce a variety of biological and technical complexities that bias relative read abundance (Fonseca, 2018; Iacaruso et al., 2025; Ruppert et al., 2019; Wu et al., 2026). However, emerging evidence suggests that sequence count data from eDNA metabarcoding may be able to track conventional survey metrics, particularly when assays are designed to amplify DNA derived from specific taxa of interest. In multispecies fish assemblages, for example, sequence counts were found to correlate with species biomass in controlled mesocosm experiments (Kelly et al., 2014), scale with abundance estimates in replicated freshwater communities (Evans et al., 2017), and reflect trawl-derived biomass, abundance, and catch per unit effort estimates from the fishing vessels (Stoeckle et al., 2021; Thomsen et al., 2016; Urban et al., 2023).

Applications to monitoring reef-building coral communities are more limited but also promising. Early work using coral-tailored 16S and CO1 markers identified a positive relationship between eDNA sequence count and live coral cover in a low-diversity Hawaiian coral community (Nichols & Marko, 2019). Subsequent studies using a tailored hypervariable internal transcribed spacer 2 (ITS2) region have offered greater resolution in several reefs of the biodiverse Indo-Pacific (Alexander et al., 2020; Dugal et al., 2022; Peiris et al., 2024; West et al., 2022), but the need to use two assays to capture the full diversity of Scleractinia complicates interpretations of relative abundance data. More recently, a 12S rDNA assay has demonstrated high taxonomic specificity and detectability in corals from complex and diverse samples (Hisata et al., 2025; Narisoko et al., 2026; Nishitsuji et al., 2023; Noda et al., 2025; Shinzato et al., 2021); however, the extent to which this assay reflects coral cover, community assemblage, and changes following severe bleaching mortality remains unclear.

During the austral summer of 2025, Mermaid Reef in remote northwest Australia experienced unprecedented thermal stress that resulted in catastrophic bleaching mortality across reefs, habitats and coral taxa. Here, we use samples collected before and after the bleaching event at Mermaid Reef to test whether coral-targeted seawater eDNA metabarcoding captures the same community change detected by conventional photo-transect surveys. Specifically, we ask whether eDNA relative read abundance tracks photo-transect percentage cover, whether the two methods capture similar genus-level shifts in community composition, and whether severe bleaching mortality is accompanied by losses in detectable coral barcode diversity.

## Methods

### Photo-transect data collection and processing

The Rowley Shoals comprise three oceanic atolls (Clerke, Imperieuse, and Mermaid Reefs), characterized by high coral cover, low nutrient inputs, and minimal local human impacts (Gilmour et al., 2019). The northernmost Mermaid Reef is a no-take Commonwealth National Park (North-west Marine Parks Network Management Plan 2018). Six long-term monitoring (LTM) sites at Mermaid Reef, spanning representative reef habitats historically dominated by *Acropora* (lagoon, crest, slope), were surveyed in October 2023, October 2024, and May 2025. During each survey, a tape was laid along three to five 50-metre permanent transects at approximately three and six metres depth, and images of the benthic community were captured at regular intervals of 1 metre from 50 cm above the substratum using a digital Canon PowerShot G1 X Mark II (12.8 MP, 1.5-inch sensor) in an underwater housing (Jonker et al., 2008). Images were analysed using a point-sampling technique to the lowest taxonomic resolution achievable. For comparison with the eDNA data, the photo-transect data were grouped to genus-level and converted to percentage coral cover (the most common metric for coral reef monitoring) for each site. This was done by averaging percent cover across replicate transects for each genus and then converting to relative coral cover (proportion of total coral cover) for each genus.

### Environmental DNA sampling, amplification, and sequencing

We collected seawater samples from the same six photo-transect LTM sites across Mermaid Reef in October 2023 and then again in May 2025. At each site, nine replicate 1-L samples were collected just beneath the surface from the side of the tender or by a snorkeller at the beginning of the permanent photo-transects. DNA from the seawater samples was concentrated on a sterile 0.45 μm cellulose nitrate membrane filter using a hand syringe and immediately preserved with DNA Shield (Zymo Research) and then stored at -20°C until processing. This resulted in three replicate filters per site, each with DNA from three litres of filtered seawater. To add confidence to our results, we also collected benthic photo-transect data and corresponding seawater samples from LTM sites across the Scott system of reefs, a neighbouring oceanic reef system 400 km to the north. In this case, samples were collected from seven LTM sites (Gilmour et al., 2019) in October 2025 after the marine heatwave using the same approach, replicates, and volumes as for the Mermaid Reef samples. Photo-transect data were also collected in parallel from the seven LTM sites using the same methodology outlined above.

DNA was then extracted from the filters using the Qiagen DNeasy Blood & Tissue Kit and the QIAcube DNA extraction robot (Qiagen) following the manufacturer’s protocol. Amplicon libraries targeting the mitochondrial 12S rRNA gene (Shinzato et al., 2021) were prepared using a two-step PCR protocol using Illumina overhang adapters (Bohmann et al., 2022). Each PCR was made up to 25 μL and contained: 12.5 µL of AmpliTaqGold 360 MasterMix, 9 µL of UltraPure Distilled water (Invitrogen), 0.5 µL of 10 μM of forward and reverse primer, 0.5 µL of GC Enhancer, and 2 µL of DNA. PCR conditions were as follows: initial denaturation at 95°C for 10 min followed by 40 cycles of 95°C for 30 s, annealing temperature at 60°C for 30 s and extension at 72°C for 45 s, with a final extension of 72°C for 10 min. Amplification was confirmed on a 2% agarose gel. All samples were processed together to minimise the impacts of batch effects, and negative controls were included throughout to check for contamination. Indexed libraries were purified, quantified, normalized, and pooled at equimolar concentrations before sequencing on an Illumina MiSeq platform using a 2 × 250 bp paired-end kit at the Australian Genome Research Facility.

### Bioinformatics and statistical analyses

Raw paired-end reads were processed in R version 4.5.0 (R Core Team) using the DADA2 package (Callahan et al., 2016), following its standard denoising workflow. Scleractinian 12S sequences were retrieved from GenBank using NCBI Entrez Direct and reformatted to create a custom reference database for taxonomic assignment in DADA2 and only amplicon sequence variants (ASVs) belonging to the order Scleractinia were retained. Rare ASVs were discarded if they occurred below a minimum sample threshold detection limit of 0.01% or detected in less than five samples.

The relationship between photo-transect and eDNA data for overlapping taxa was first tested using a Gaussian general linear model, with photo-transect relative cover (proportion of total coral cover) as the response variable and eDNA relative read abundance (proportion of total reads) as the predictor. Site, year and genus were included as fixed effects to account for spatial, temporal and taxonomic differences among observations. Pearson correlation was also calculated on the relative abundance and relative cover values to provide an unadjusted measure of association between methods. For samples from Scott Reef, we used the same modelling framework as for Mermaid Reef, except for omitting sample year. Community-level consistencies between methods were assessed using the Mermaid site-year by genus matrices, where relative abundance data were square-root transformed and used to estimate Bray-Curtis dissimilarity matrices for each method. These matrices were then compared using a Mantel test based on 9,999 permutations. Agreement between the community distance matrices was also assessed using two-dimensional NMDS ordinations using Procrustes analysis (9,999 permutations). To assess whether this relationship was robust to bioinformatic filtering, we repeated the Mantel tests across a range of ASV filtering thresholds, where filtering combinations included minimum depth, prevalence, minimum relative abundance, minimum sample read depth, and the threshold used to set low abundance reads to zero. For each filtering scenario, non-coral ASVs, negative controls, and low-depth samples were removed, and remaining ASVs were collapsed to genus level, and then Bray-Curtis dissimilarities were recalculated and retested using Mantel tests. Finally, we tested whether bleaching altered coral barcode diversity, measured as ASV richness, at both the whole-site and genus levels. Site-level ASV richness was compared before and after bleaching using a paired t-test, while genus level changes were tested using a mixed-effects model with sample year as the fixed effect and site and genus as random effects.

## Results and Discussion

Coral cover at Mermaid Reef in October 2023 was high, ranging from 24–49% across the six long-term monitoring sites (Table S1). The diverse coral communities comprised 29 different genera, with approximately half of the mean coral cover belonging to *Acropora* (17% ± 11.34 SD). *Montipora* and *Porites* were also common across the reef, though each comprised less than 5% of total coral cover. In October 2024, prior to the onset of heat stress, photo-transect surveys revealed healthy coral communities with high coral cover similar to the previous year (Figure 1b, Table S1). However, by February 2025, temperatures at Mermaid Reef had already exceeded 12 Degree Heating Weeks (DHW), far exceeding established bleaching thresholds (Figure 1a). As a result, the reef suffered its most severe bleaching mortality event on record, and from October 2024 to May 2025, reef-wide hard coral cover declined from 35.3% (± 12.4 SD) to 5.9% (± 3.74 SD) across our LTM sites (Figure 1b). *Acropora* was the most impacted and declined to less than 0.5% cover across all sites (Figure 1c). Other common taxa, like *Porites* and *Montipora*, declined to a lesser extent but each accounted for less than 1% cover after bleaching.

**Figure 1.**
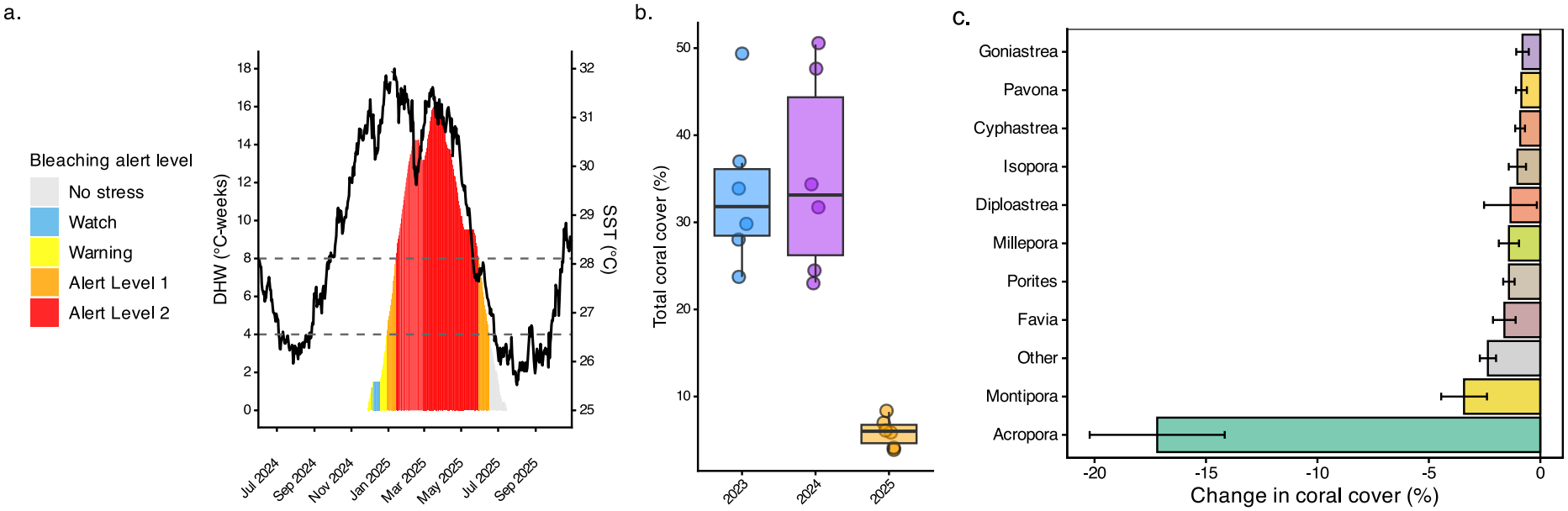
Severe heat stress and coral mortality at Mermaid Reef: a) NOAA Coral Reef Watch virtual stations data showing accumulation of heat stress at the Rowley Shoals during the 2024/25 austral summer; b) estimates of coral cover from diver photo-transects at long-term monitoring sites prior to heat stress (October 2023, 2024), and after heat stress in May 2025. Each point represents a site mean averaged across all transects. Interquartile ranges are plotted behind points; c) absolute change (± SE) in percentage coral cover following mass bleaching in the top 10 most abundant genera of coral.

This severe decline in *Acropora* and the shift in relative abundance of other taxa was detected by amplifying and sequencing coral DNA from seawater. Amplicon sequencing of the 12S rRNA region produced 3,877,636 raw paired end reads that clustered into 2,819 unique amplicon sequence variants (ASVs; Table S2), 144 of which could not be classified as Scleractinia and were discarded (Table S2). After quality control and filtering, 456 ASVs in 38 samples remained (Figure S2). In this dataset we recovered ASVs from 20 different genera of scleractinian coral (Figure 2a, b; Table S3), 15 of which overlapped with genera present in the photo-transect data (Figure 2c). The ASVs belonging to the shared 15 genera accounted for 95.8% of the total reads recovered from seawater samples and 86% of the total coral cover from photo-transect data. These data revealed that pre-bleaching samples were strongly dominated by sequence counts from the genus *Acropora*, accounting for 78% of all reads and comprising 65 unique ASVs (Figure 2a). *Porites* (8.9% of reads and 56 ASVs) and *Montipora* (4.32% of reads and 31 ASVs) were the next most abundant taxa. Samples collected in May 2025 following bleaching mortality showed a strong decline in *Acropora* sequence counts. *Acropora* relative read abundance declined by 63% (Figure S3), and less abundant taxa that were previously detected at levels below 10%, like *Porites* and *Montipora*, increased to 32% and 21%, respectively (Figure 2d).

**Figure 2.**
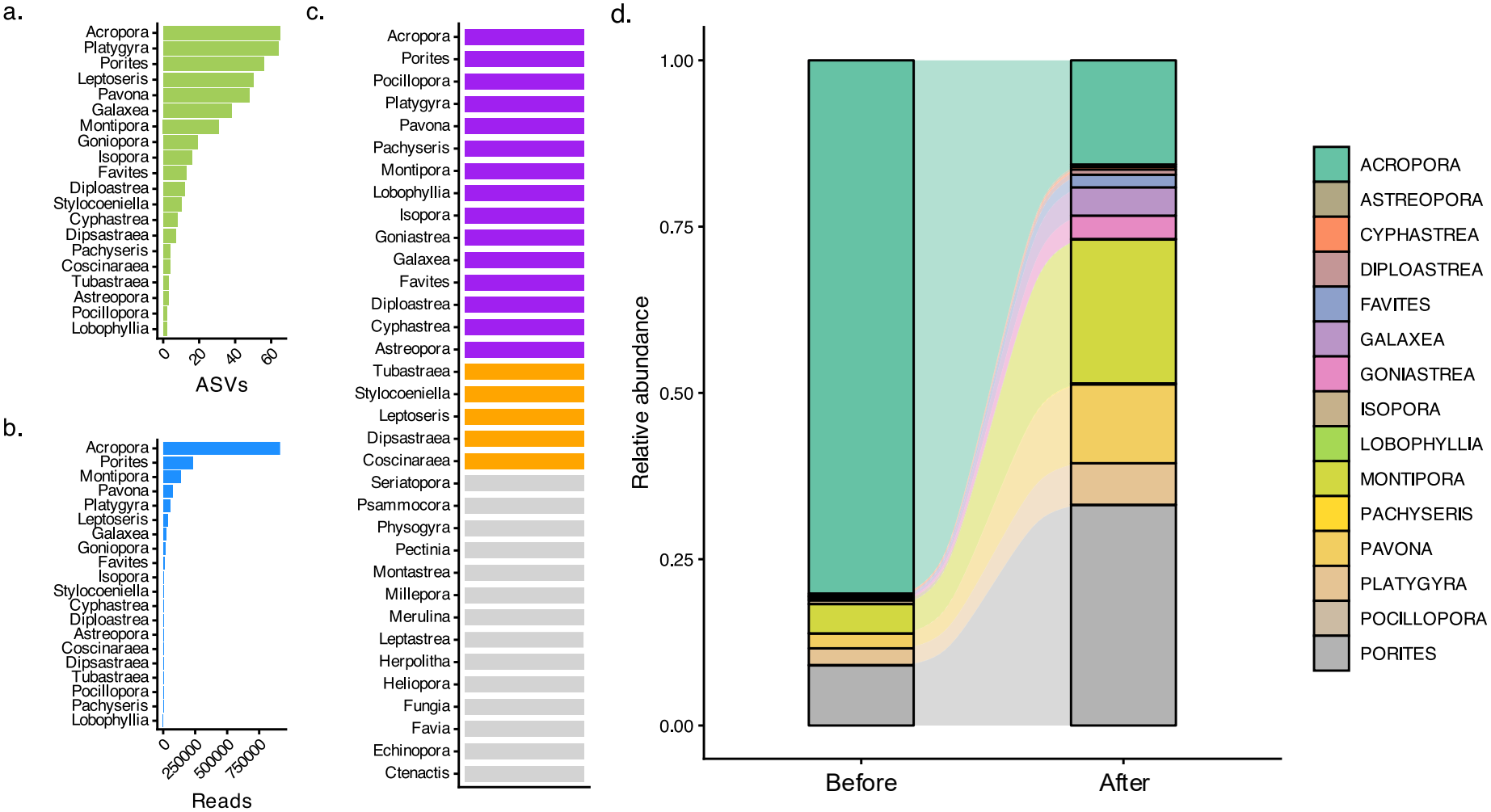
Monitoring changes to coral communities with seawater eDNA metabarcoding: a) bar plots of ASV diversity and b) sequence counts for each taxon recovered from eDNA metabarcoding; c) tile plot showing genera recovered by the two methods (n = 15, purple), unique to eDNA (n=5, orange) and unique to photo-transect data (n=14, grey); d) alluvial plot showing shift in eDNA relative abundance before and after the heat stress.

Across sites and time points, photo-transect relative coral cover was strongly associated with eDNA relative read abundance for the 15 coral genera detected by both methods (Pearson’s r = 0.85, P < 0.001; Figure 3a, Figure S4). This positive association was supported by a Gaussian general linear model in which photo-transect-derived relative cover increased with eDNA relative read abundance after accounting for site, sample year, and genus identity (β = 0.700, P < 0.001). Community-level patterns were also consistent between methods. Mantel tests showed a significant relationship between Bray-Curtis dissimilarity matrices derived from photo-transect and eDNA relative abundance data (r = 0.750, P < 0.001; Figure 3b) that was robust to varying sample and ASV filtering thresholds (Figure S5). Furthermore, Procrustes analysis of NMDS ordinations showed strong alignment between estimates of community structure (r = 0.770, P = 0.002; Figure 3c). Finally, an independent sampling and sequencing run from Scott Reef in October 2025 showed the same positive association between eDNA relative read abundance and relative coral cover, adding support to our findings (Pearson’s r = 0.657, P < 0.001; Figure S6, S7, Table S5, S6, S7).

**Figure 3.**
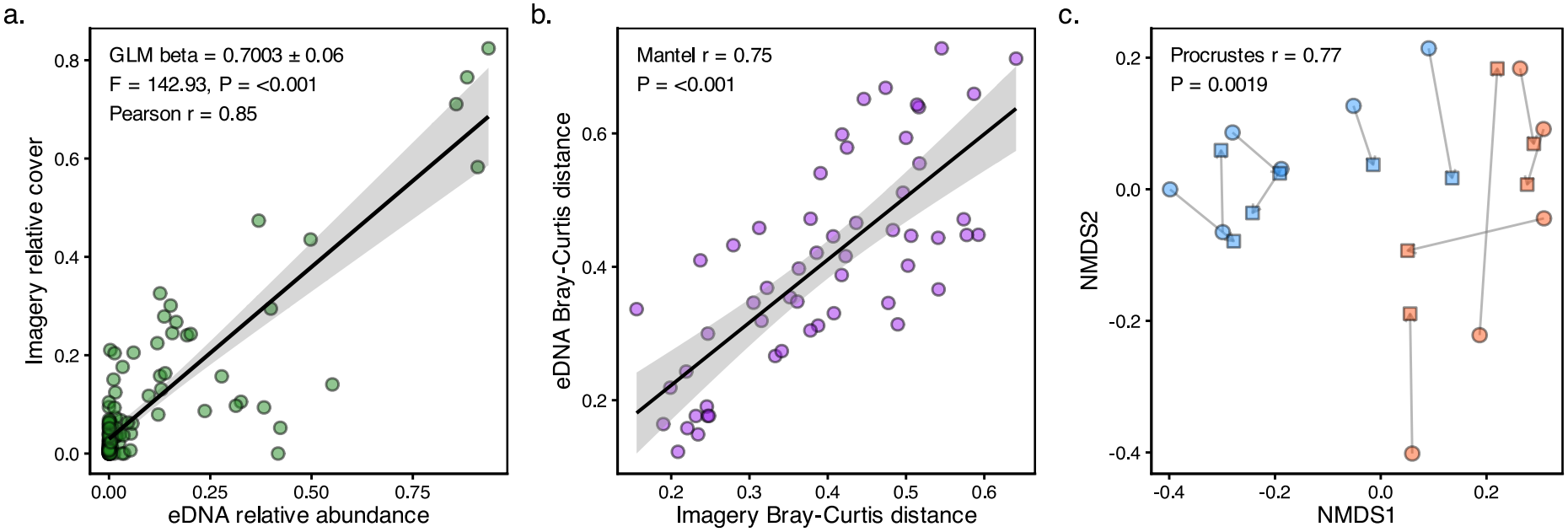
Relationship between eDNA and photo-transect derived metrics of relative coral cover: a) Modelled relationship between eDNA relative read abundance and photo-transect derived relative coral cover (proportion of total coral cover) across the 15 overlapping coral taxa and long-term monitoring sites. Points represent genus-by-site observations, and the fitted line shows the estimated relationship from the general linear model; b) Relationship between Bray-Curtis dissimilarity matrices calculated from eDNA and photo-transect data. Each point represents a pairwise site comparison; c) Procrustes ordination comparing genus level community structure inferred from eDNA relative read abundance (squares) and photo-transect relative cover (circles) across matched sites before (blue) and after (red) bleaching. Lines connect paired eDNA and photo-transect ordinations for the same site and timepoint, with shorter lines indicating stronger agreement between methods.

The decline in coral cover at Mermaid Reef was not accompanied by a consistent loss of total coral barcode diversity. Total coral ASV richness per site remained similar before and after bleaching (Figure 4a). However, genus level ASV richness declined following bleaching (β = ™4.78, P = 0.002). This decline was taxon-specific, with the largest declines in *Acropora* and *Leptoseris*, while other genera showed mixed or non-significant changes (Figure 4b, 4c). These results suggest that bleaching did not cause an immediate loss of detectable coral barcode richness at the reef scale, but instead reduced barcode richness within certain taxa and changed how barcode diversity was distributed among sites. Disturbances often restructure ecological communities before producing measurable losses of diversity (Dornelas et al., 2014; Hillebrand et al., 2018; Richards et al., 2021). Here, the strongest eDNA signal was not a wholesale loss of coral barcode richness, but a redistribution of relative read abundance and ASV richness among less abundant surviving genera.

**Figure 4.**
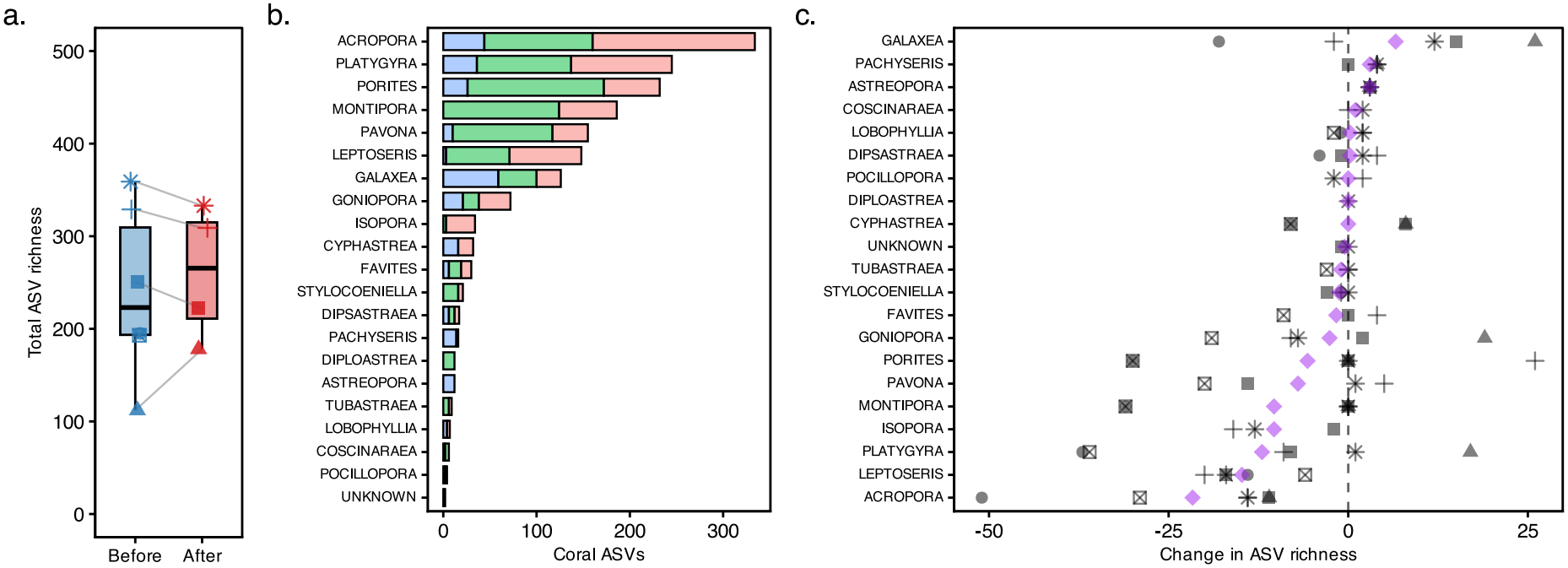
Changes in barcode diversity in response to bleaching mortality: a) Boxplots of ASV richness across all coral taxa before and after bleaching mortality. Symbols represent sites connected by grey lines; b) bar plot of coral ASVs classified as lost (red), retained (green), or gained (blue) between pre and post-bleaching surveys; c) change in ASV richness after bleaching for each genus. Purple diamonds indicate mean difference across sites and shapes represent different sites.

Recent studies show that coral-targeted metabarcoding can recover broad diversity patterns, distinguish reef assemblages, and in some cases reflect visual estimates of coral cover (Alexander et al., 2020; Dugal et al., 2022; Hisata et al., 2025; Nichols & Marko, 2019; Nishitsuji et al., 2023; Peiris et al., 2024; Shinzato et al., 2021). We extend this work by demonstrating that metabarcoding can track temporal changes in relative coral cover and capture major compositional shifts following severe mortality. Photo-transects yield direct estimates of live cover and eDNA provides a scalable, taxonomically sensitive complement to track shifts in dominance, barcode richness, and genus level diversity through time. After severe disturbances, eDNA may detect low-abundance taxa missed by photo-transects, although these detections may derive from surviving or cryptic colonies, transported DNA, or recently dead tissue, and therefore do not directly indicate live coral cover. Where detections reflect remnant live colonies, their low density is ecologically important because spatial isolation can limit fertilisation and recruitment through Allee effects (Mumby et al., 2024; Ricardo et al., 2026). Taken together, our findings support the broader adoption of eDNA sampling as a complementary component of coral reef monitoring.

## Acknowledgements

This dataset was collected during multiple research cruises, led by the Australian Institute of Marine Science (AIMS), as part of the Ocean Discovery & Restoration Program and supported by Parks Australia.

---

**Table S1.** Absolute coral cover (mean across transects and depths (%)) at each site based on photo-transect data collected by divers on SCUBA.

| **Site** | **October 2023** | **October 2024** | **May 2025** |
|---|---|---|---|
| **M11** | 28.06 | 23.09 | 3.80 |
| **M13** | 23.73 | 24.37 | 6.93 |
| **RL1-N** | 33.94 | 34.50 | 5.79 |
| **RL1-S** | 36.83 | 50.42 | 4.25 |
| **RS1-1** | 29.67 | 31.78 | 6.23 |
| **RS1-S** | 49.21 | 47.65 | 8.23 |

**Table S2.** Sequencing and clustering results for the Mermaid Reef eDNA dataset collected before and after bleaching mortality, including number of samples, total number of amplicon sequence variants matching Scleractinia, total reads that passed through QC, mean reads per sample, and mean ASVs per sample.

|  | **Samples** | **ASVs** | **Total reads** | **Mean reads** | **Mean ASVs** |
|---|---|---|---|---|---|
| **DADA2** | 42 | 2,819 | 1,689,883 | 40,235 | 228 |
| **ASV filtering** | 42 | 456 | 1,523,453 | 36,273 | 158 |
| **Sample filtering** | 38 | 456 | 1,510,486 | 39,750 | 170 |

**Table S3.** Taxonomic classification of amplicon sequence variants from seawater DNA. In total, we recovered 20 unique genera from Mermaid Reef.

| **Genus** | **ASVs** | **Total reads** | **Amplicon length (bp)** |
|---|---|---|---|
| **Acropora** | 65 | 914110 | 453 |
| **Porites** | 56 | 231723 | 434 |
| **Montipora** | 31 | 135229 | 453 |
| **Pavona** | 48 | 72285 | 453 |
| **Platygyra** | 64 | 52271 | 368 |
| **Leptoseris** | 50 | 33053 | 445 |
| **Galaxea** | 38 | 20250 | 453 |
| **Goniopora** | 19 | 16844 | 433 |
| **Favites** | 13 | 8894 | 368 |
| **Isopora** | 16 | 5866 | 453 |
| **Stylocoeniella** | 10 | 4549 | 447 |
| **Cyphastrea** | 8 | 4220 | 368 |
| **Diploastrea** | 12 | 3798 | 367 |
| **Astreopora** | 3 | 1476 | 453 |
| **Coscinaraea** | 4 | 1452 | 371 |
| **Dipsastraea** | 7 | 1414 | 369 |
| **Tubastraea** | 3 | 826 | 434 |
| **Pocillopora** | 2 | 800 | 379 |
| **Pachyseris** | 4 | 727 | 453 |
| **Lobophyllia** | 2 | 452 | 367 |

**Table S4.** Relationship between eDNA relative abundance and photo-transect-derived relative coral cover (proportion of total) at Mermaid Reef.

| **Statistic** | **Estimate** | **P value** |
|---|---|---|
| **Pearson r** | 0.848 | <0.001 |
| **GLM Beta** | 0.700 | <0.001 |
| **Mantel r** | 0.750 | <0.001 |
| **Procrustes r** | 0.770 | 0.002 |

**Table S5.** Sequencing statistics for samples of seawater DNA collected from Scott Reef.

| **Samples** | **ASVs** | **Reads** | **Sites** |
|---|---|---|---|
| 26 | 286 | 2340177 | 7 |

**Table S6.** Taxonomic classification of amplicon sequence variants from seawater DNA collected from Scott Reef. In total, we recovered 15 unique genera from Scott Reef, 11 of which overlapped with the photo-transect data.

| **Genus** | **ASVs** | **Total Reads** |
|---|---|---|
| **Porites** | 43 | 1136508 |
| **Acropora** | 27 | 720877 |
| **Montipora** | 15 | 167485 |
| **Leptoseris** | 19 | 84193 |
| **Pavona** | 29 | 75802 |
| **Platygyra** | 61 | 62380 |
| **Stylocoeniella** | 26 | 36682 |
| **Isopora** | 16 | 18095 |
| **Galaxea** | 18 | 8480 |
| **Pocillopora** | 8 | 4370 |
| ***Letepsammia*** | *15* | *3623* |
| ***Stephanocyathus*** | *3* | *716* |
| ***Deltocyathus*** | *2* | *668* |
| ***Coscinaraea*** | *2* | *646* |
| ***Stephanocoenia*** | *2* | *522* |

**Table S7.** Relationship between eDNA relative abundance and photo-transect derived relative coral cover at Scott Reef.

| **Statistic** | **Estimate** | **P value** |
|---|---|---|
| **Beta** | 0.079 | 0.006 |
| **Pearson r** | 0.657 | <0.001 |

**Figure S1.**
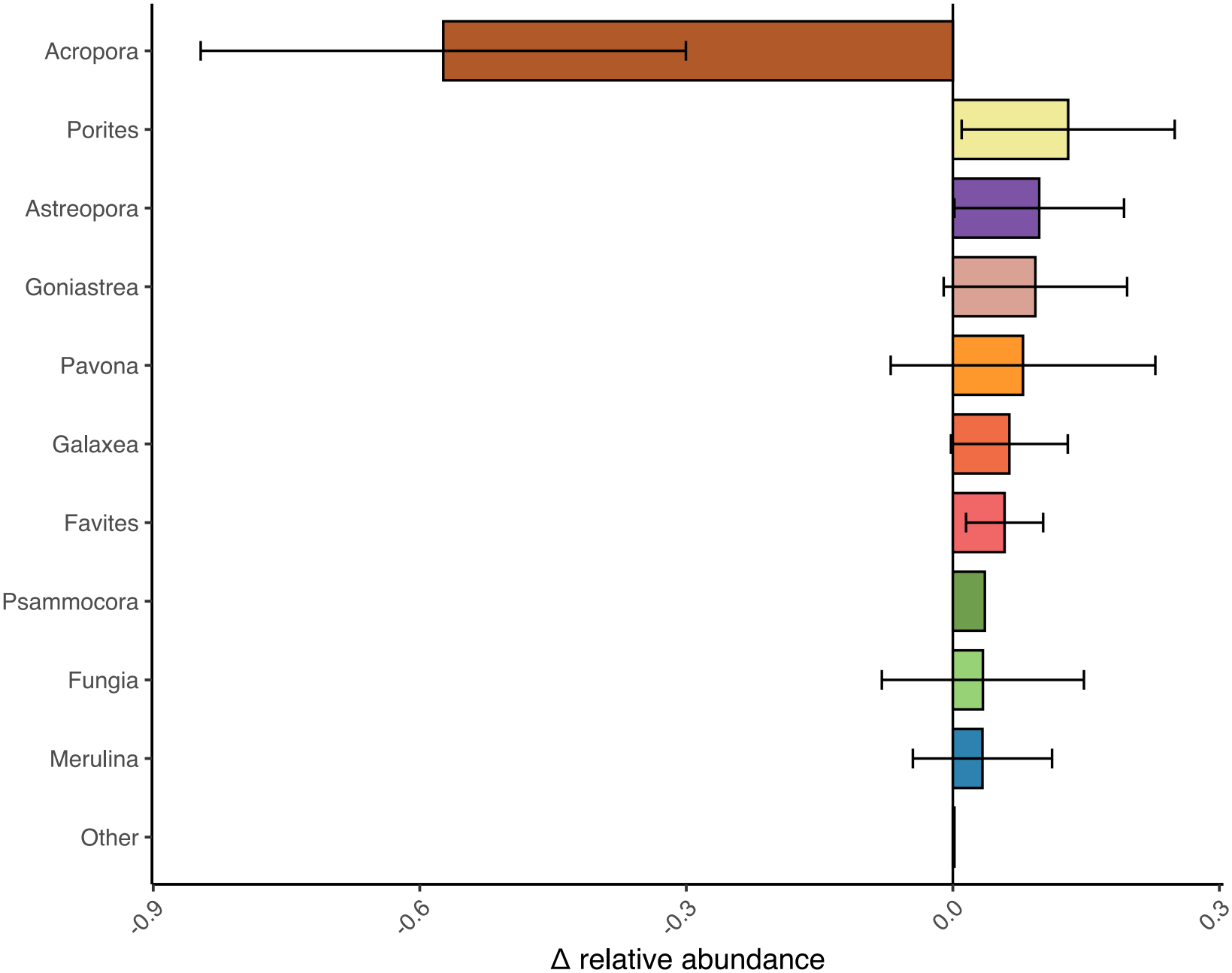
Horizontal bar plots of relative abundance (with standard error) for the dominant coral genera based on photo-transect data following bleaching mortality at Mermaid Reef.

**Figure S2.**
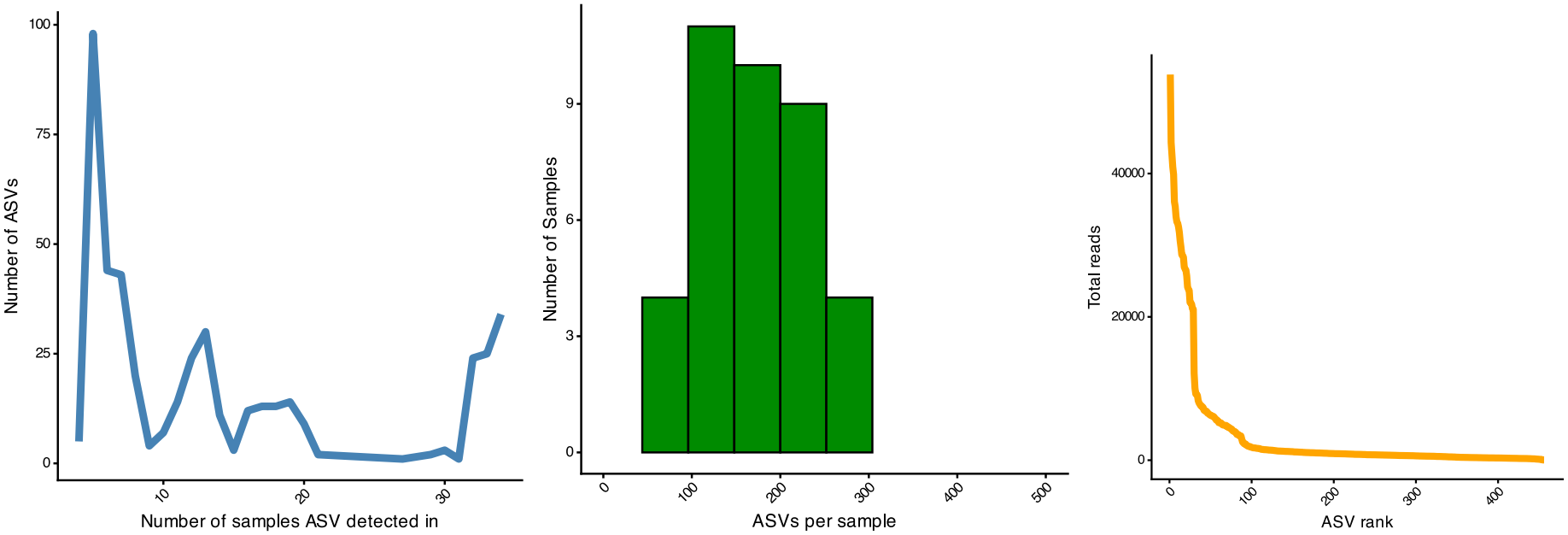
Summary statistics of the eDNA ASV dataset, including the frequency of ASV detection across samples, the distribution of ASV richness per sample, and total read abundance per ASV.

**Figure S3.**
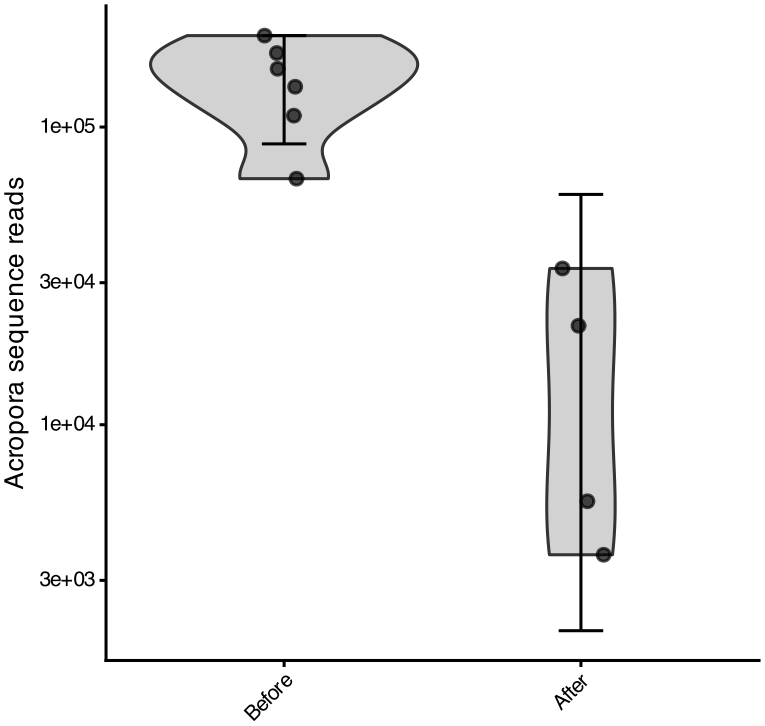
Acropora sequence reads recovered from seawater samples collected before and after bleaching mortality.

**Figure S4.**
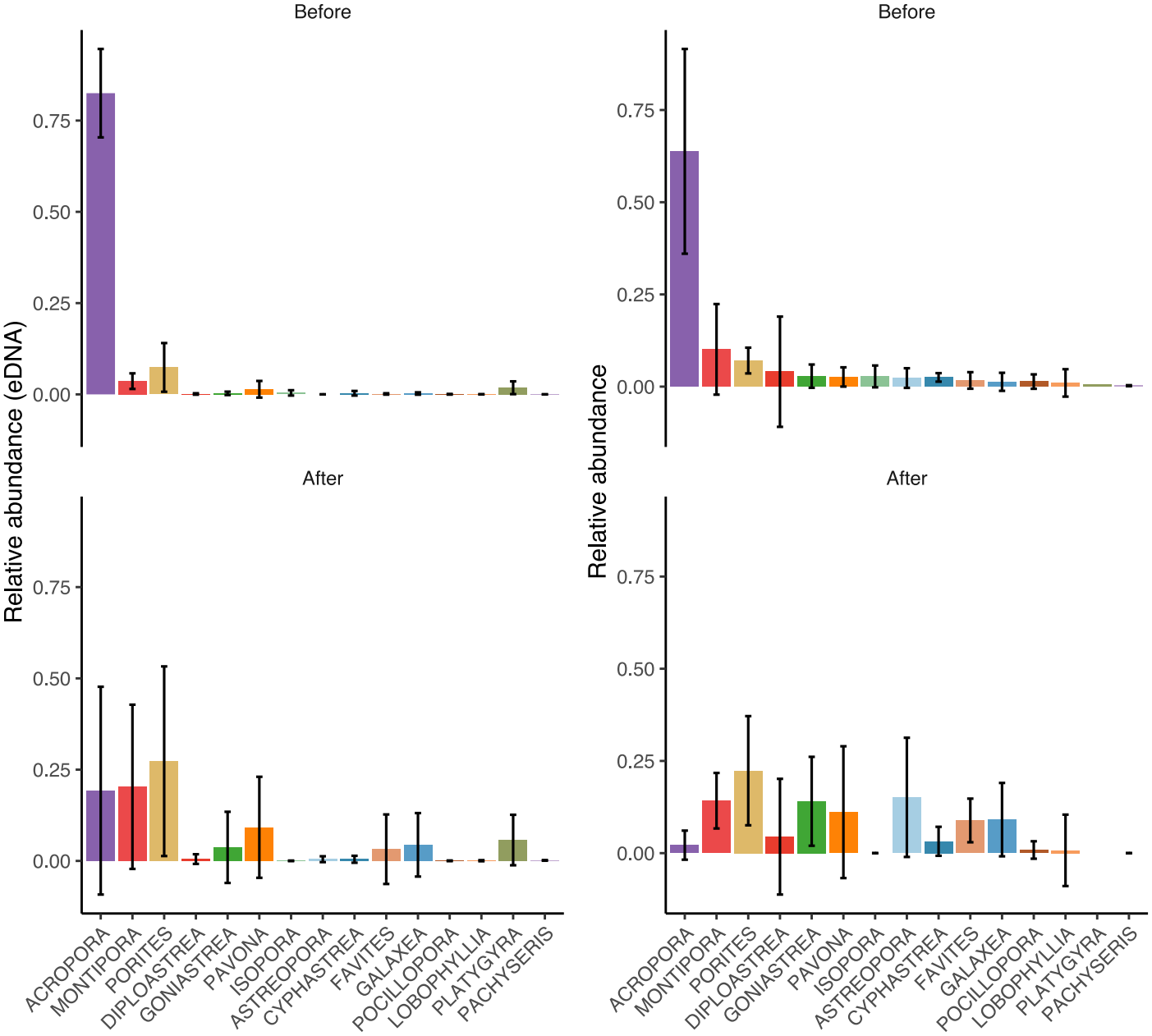
Bar plots of mean relative abundance (± SE) of the 15 coral genera shared between photo-transect surveys and eDNA metabarcoding, shown for each method across the two sampling timepoints.

**Figure S5.**
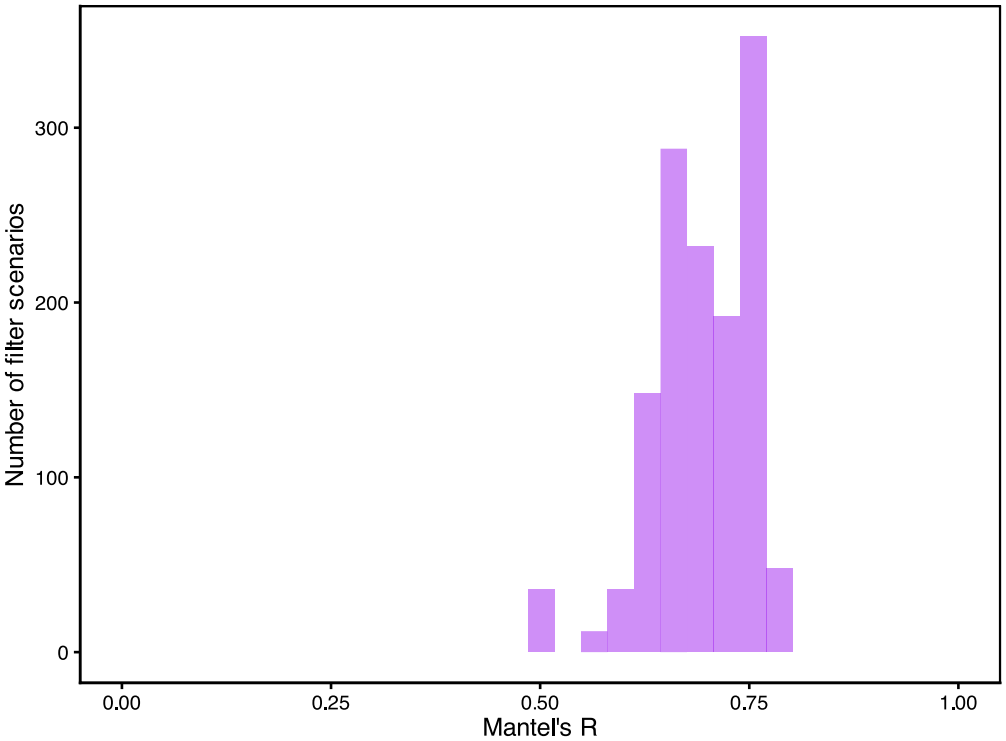
Histogram of Mantel’s r based on filter threshold sensitivity analysis comparing eDNA relative read abundance and photo-transect derived relative coral cover.

**Figure S6.**
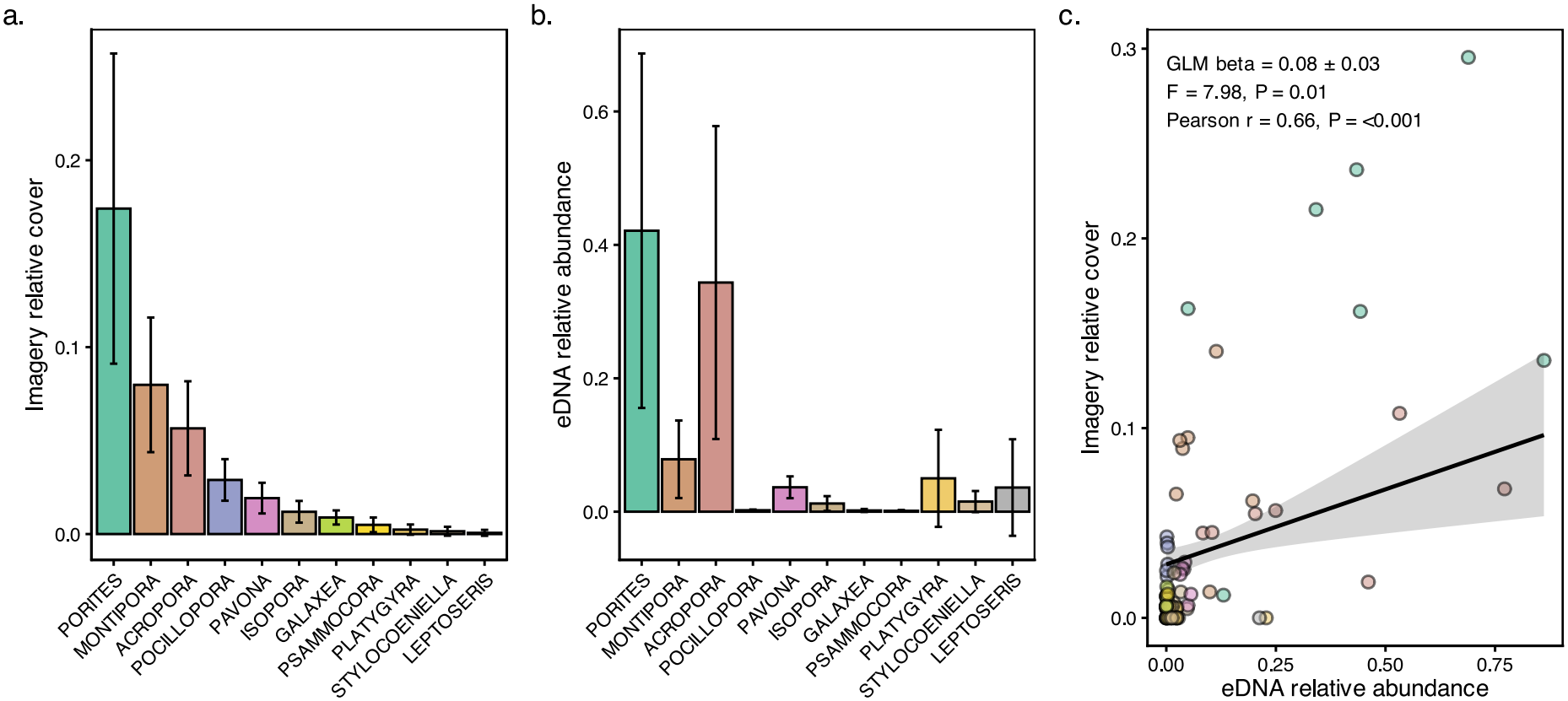
Bar plots of mean relative abundance (± SE) of the 11 coral genera shared between photo-transect surveys (a) and eDNA metabarcoding (b) at Scott Reef. Relationship between eDNA relative read abundance and photo-transect-derived relative coral cover across Scott Reef sites (c). Points represent site-by-genus observations. The black line shows the fitted Gaussian GLM relationship, with the shaded band representing the 95% confidence interval. Photo-transect-derived relative cover was positively associated with eDNA relative read abundance after accounting for site and genus identity (β = 0.079 ± 0.028 SE, F = 7.98, P = 0.006). The unadjusted Pearson correlation was also positive (r = 0.66, P < 0.001).

**Figure S7.**
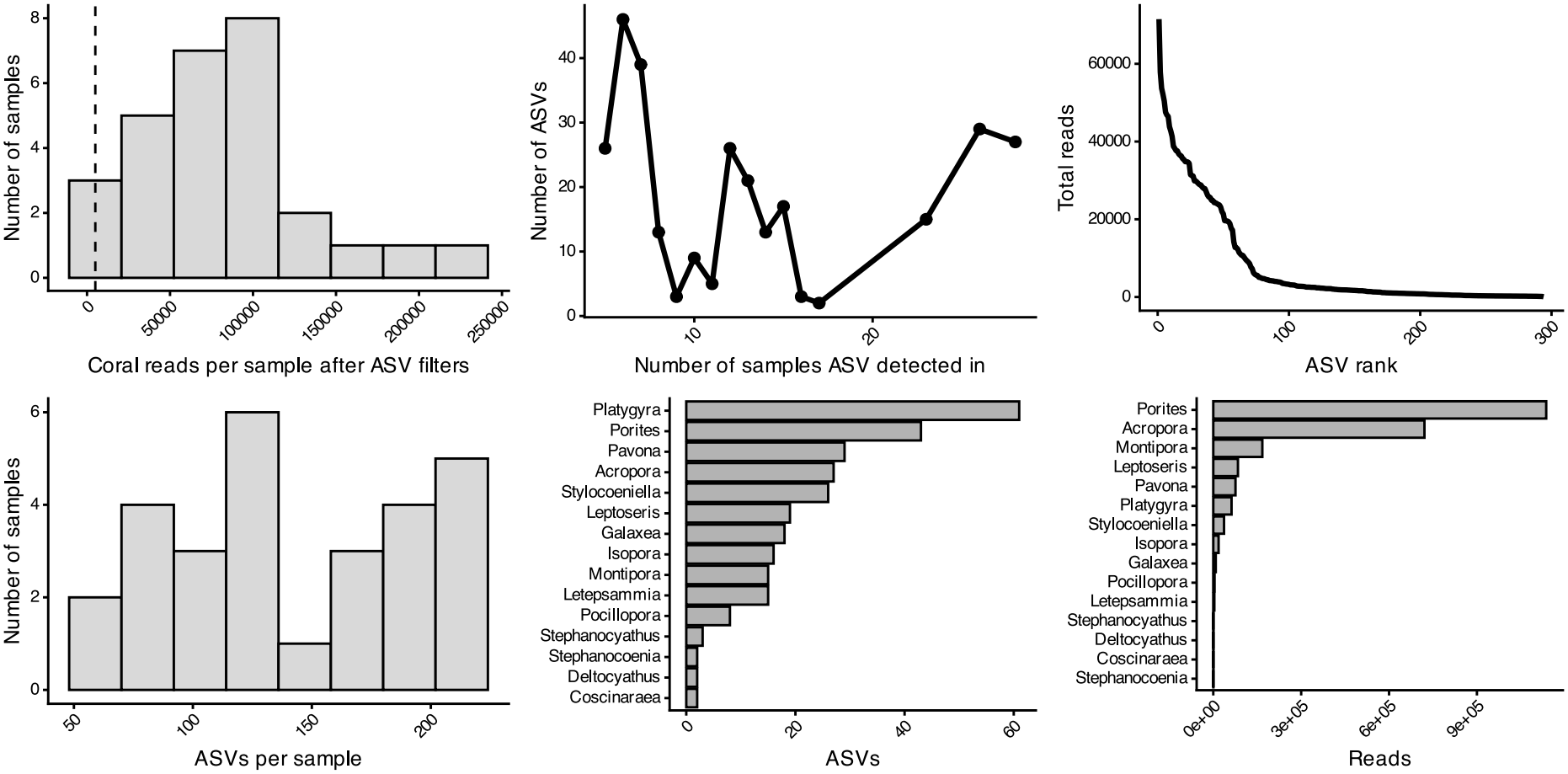
Additional sequencing results and taxonomic output from Scott Reef samples. Top plots show histogram of reads per sample (left), number of ASVs found across samples (middle) and total reads per ASV (right). Bottom plots show histogram of ASVs per sample (left), number of ASVs (middle) and reads (right) per taxa.

**Figure S8.**
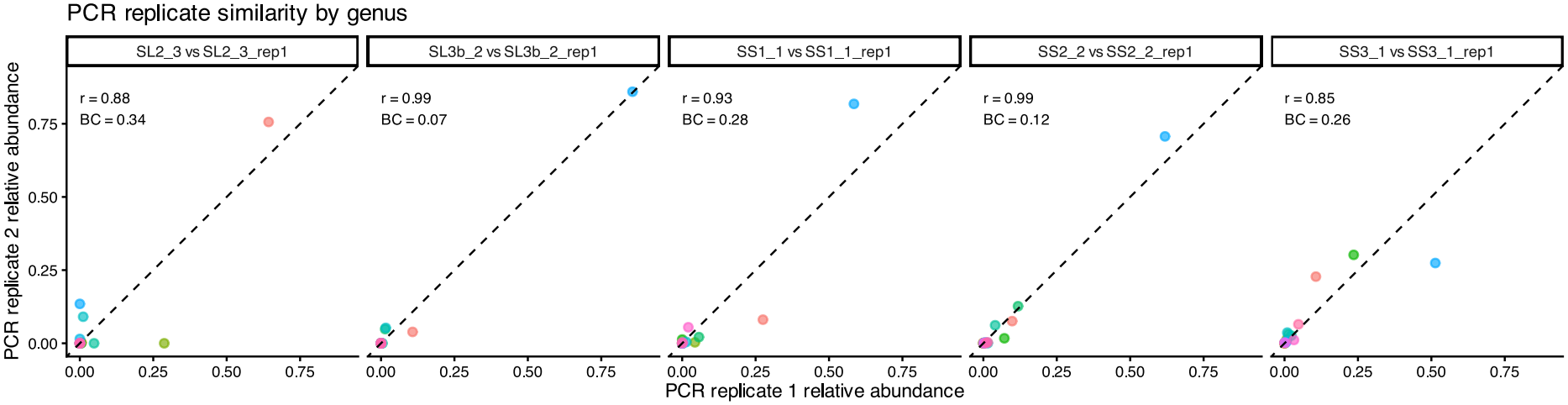
Reproducibility of relative abundance between PCR replicates; scatter plots compare genus (coloured points) relative abundance between replicate PCR libraries, with the dashed line indicating a 1:1 relationship. Pearson correlation and Bray-Curtis dissimilarity are shown for each replicate pair.

